# Losses resulting from deliberate exploration trigger beta oscillations in frontal cortex

**DOI:** 10.1101/2022.10.06.510948

**Authors:** Boris V. Chernyshev, Kristina I. Pultsina, Vera D. Tretyakova, Aleksandra S. Miasnikova, Andrey O. Prokofyev, Galina L. Kozunova, Tatiana A. Stroganova

**Affiliations:** Center for Neurocognitive Research (MEG Center), Moscow State University of Psychology and Education, Moscow, Russia; Department of Higher Nervous Activity, Lomonosov Moscow State University, Russia; Department of Psychology, Higher School of Economics, Moscow, Russia

**Keywords:** decision making, exploration, exploration-exploitation dilemma, gambling, probabilistic task, magnetoencephalography

## Abstract

We examined the neural signature of directed exploration by contrasting MEG beta (16-30 Hz) power changes between disadvantageous and advantageous choices in the two-choice probabilistic reward task. We analyzed the choices made after the participants have learned the probabilistic contingency between choices and their outcomes, i.e., acquired the inner model of choice values. Therefore, rare disadvantageous choices might serve explorative, environment-probing purposes. The study brought two main findings. Firstly, decision making leading to disadvantageous choices took more time and evidenced greater large-scale suppression of beta oscillations than its advantageous alternative. Additional neural resources recruited during disadvantageous decisions strongly suggest their deliberately explorative nature. Secondly, an outcome of disadvantageous and advantageous choices had qualitatively different impact on feedback-related beta oscillations. After the disadvantageous choices, only losses – but not gains – were followed by late beta synchronization in frontal cortex. Our results are consistent with the role of frontal beta oscillations in the stabilization of neural representations for selected behavioral rule when explorative strategy conflicts with value-based behavior. Punishment for explorative choice being congruent with its low value in the reward history is more likely to strengthen, through punishment-related beta oscillations, the representation of exploitative choices consistent with the inner utility model.

## 1 Introduction

In professional and everyday life – for example, in search of a new job – people often face an exploration-exploitation dilemma. Searching for a new opportunity at the expense of the familiar ones can be costly, as tangible outcomes of exploration may only come in the distant future and no profits are warranted. On the other hand, sticking to the choices that proved valuable in the past discourages subjects from pursuing learning and development. A purposeful search for information at the expense of immediate reward is a characteristic feature of directed exploration, which became topic of considerable interest the last decade (Payzan-LeNestour and Bossaerts, 2012; Wilson et al., 2014; Zajkowski et al., 2017; Schulz and Gershman, 2019; Schwartenbeck et al., 2019; Dubois et al., 2021; Wilson et al., 2021). Although switching between exploitative and deliberate explorative strategies constitutes a core foundation of human adaptive behavior, its neural underpinnings remain largely unknown (but see Zajkowski et al., 2017; Gottlieb and Oudeyer, 2018; Koechlin, 2020).

People actively interrogate their environment using ‘question-and-answer’ strategies (Gottlieb and Oudeyer, 2018). Yet this is not an easy challenge: choosing a risky and potentially non-rewarding option involves a conflict, since participants have to inhibit the tendency to choose a rewarded safe alternative (Daw et al., 2006; Cogliati Dezza et al., 2017). This conflict arises between at least two simultaneously active competing internal models, or ‘task sets’ (Domenech et al., 2020; Koechlin, 2020) – one being a predominant response tendency (exploitation), and the other – its conscious alternative (exploration). Our recent pupillometric study lends support to this assumption (Kozunova et al., 2022): we found that such explorative choices compared to exploitative ones are accompanied by larger pupil dilation and longer decision time. We speculated that this state of conflict supposedly entails an increase in the degree of processing required to make the deliberately explorative decisions. Given that such a selection was shown to carry greater load on information processing and/or attentional capacities (Gottlieb and Oudeyer, 2018), it should make a deliberately explorative decision more difficult than the exploitative decision. To validate this explanation, one needs to compare cortical activation accompanying the decision process that leads to either exploitative or to explorative choices.

Importantly, any action outcome – even a negative one – has epistemic value since it reduces internal uncertainty (Parr and Friston, 2017). Since deliberate explorative choices are in fact quests for information, they likely involve altered evaluation of the outcome received after one’s choice. Indeed, for example, even modest variations in the context – such as changing the subject’s beliefs about the probability of rewards and punishments – can alter cortical responses to feedback signals (e.g., Marco-Pallares et al. (2008)).

Here, we sought to evaluate the impact of directed exploration on cortical signatures of decision-making and feedback evaluation processes. To this end, we investigated the dynamics of magnetoencephalographic β-band oscillations (16-30 Hz) while contrasting directed explorative choices and exploitative ones in the probabilistic two-alternative task. Well-documented properties of β oscillations make them an informative and accessible measure allowing us to pursue the above goals.

Based on numerous findings on β power suppression (event-related desynchronization, β-ERD) as a measure of cortical activation strength during decision-making (Scharinger et al., 2017; Pavlova et al., 2019; Tafuro et al., 2019), we predicted that decision-making leading to directed explorative choices of the apparently disadvantageous option would be accompanied by greater β-ERD than decision-making resulting in exploitative choice.

Concerning β-oscillations over the frontal cortex (frontal event-related synchronization, frontal β-ERS) following action outcome evaluation, the previous literature offers two accounts. The first account relies on the evidence specifically linking this oscillatory activity with reward, especially with an unexpected reward in human subjects (Marco-Pallares et al., 2008; Mas-Herrero et al., 2015). This view is supported by the findings on greater frontal β-ERS for positive compared to negative feedback (Cohen et al., 2011; Weiss and Mueller, 2012; Novikov et al., 2017) and for larger compared to smaller reward magnitudes (Marco-Pallares et al., 2008). It was proposed that feedback-related frontal β-ERS reflects a signal involved in reward processing and underlying memory formation by signaling which events are better than expected (for review, see Marco-Pallares et al. (2015). Based on the ‘reward account’ one can predict that in our study, a reward will be followed by frontal β-ERS compared to punishment regardless of the type of the choice and will be relatively greater for explorative compared with exploitative choices – simply because for explorative choices a positive reward is less expected.

Still, the dependence of frontal β-oscillations on some specific task conditions is difficult to explain using the straightforward ‘reward account’. For example, there is evidence that in some tasks, a negative feedback signal (or omission of a positive one) contributes to the build-up of the inner state promoting the increase in frontal β-oscillations (Leicht et al., 2013; Yaple et al., 2018).

A complementary account, which accumulates numerous findings on frontal β-synchronization in both humans and animals, proposes that these oscillations play a role in re-activation and strengthening of the cognitive set, which is more likely to result in a favorable outcome and will be used in future actions (Engel and Fries, 2010; Brincat and Miller, 2016; Miller et al., 2018). The term ‘cognitive set’ here is very similar to the term ‘internal model’, which, according to K. Friston’s predictive coding theory, accumulates experience-based knowledge and generates prior beliefs that are used to make inferences/predictions regarding incoming external events (Friston and Kiebel, 2009; Summerfield and de Lange, 2014). From a predictive coding perspective, for the selection of a to-be-strengthened cognitive set/internal model, both positive and negative outcomes may be used as just pieces of evidence weighted against recent reward history that formed the inner utility model (Kennerley et al., 2006; Yon and Frith, 2021). Depending on how reliable these different sources of information are, the selection process may lend the greatest weight to the inner model that ensures optimal guidance of future actions.

The ‘predictive coding’ account produces predictions different from those expected on the basis of the ‘reward’ account. In this case, appearance of frontal β-ERS will depend on how the two competing models, that were simultaneously active during decision-making regarding this choice, will be re-evaluated after receiving feedback. A reward for a rare, explorative, deliberately disadvantageous choice, which is discrepant with the previous reward history, hardly provides a piece of evidence reliable enough to ensure strengthening of the model governing this choice. Punishment for such a choice, on the other hand, being congruent with its low value in the reward history, is more likely to strengthen the representation of its competitor – the predominant utility model. This might result in the occurrence of frontal β-ERS after a punishment but not a reward for an explorative choice. In other words, directed explorative choices may be expected to differ from exploitative ones by a weak impact of a positive outcome, and a strong impact of a negative outcome.

Therefore, by evaluating the predictions of the two accounts regarding frontal β-ERS in deliberate explorative and exploitative choices, we hoped to get a better insight into the nature of brain processes resulting from the appraisal of feedback information during switching from exploitation to exploration strategy.

To sum up, we here hypothesize that disadvantageous choices in the two-alternative probabilistic reward task represent a deliberate switching from the exploitative to the directed explorative strategy that triggers a conflict with a prepotent response tendency. The modulation of brain activity in the β-range could exhibit specific features during both decision-making and feedback-related processes associated with such explorative choices. First, we sought to test the prediction that a putatively more effortful arrival to the explorative decision would involve greater cortical activation indexed by deeper β-ERD compared with exploitative decisions. Second, by contrasting post-feedback modulations of β-oscillations for the explorative and exploitative choices, we aimed to test the predictions made based on the two accounts of how prior knowledge about the probability of reward and punishment influences feedback processing in the brain.

## 2 Materials and Methods

### 2.1 Participants

Sixty healthy participants (30 males, 30 females), aged 25.6±4.8 years (M ± SD) and having no neurological disorders or severe visual impairments were enrolled in the study. This study was conducted in accordance with the Declaration of Helsinki; the study protocol was approved by the Ethics Committee of Moscow State University of Psychology and Education. All participants signed written informed consent before the experiment.

### 2.2 Procedure

During the experiment, participants were seated in a comfortable armchair with a projection screen located in front of them at eye level. A modified probabilistic learning task (Frank et al., 2004; Kozunova et al., 2018) was rendered as a computer game consisting of six blocks.

Each pair of visual stimuli comprised two images of the same Hiragana hieroglyph (1.54 × 1.44°) rotated at two different angles (Figure 1A). We equalized the stimuli in size, brightness, perceptual complexity, and spatial position. The two stimuli were placed symmetrically on the left and on the right sides of the screen 1.5° from the screen center. In order to avoid any biases related to the side of the presentation, it was swapped quasirandomly over the course of trials. In order to minimize the amount of light falling into participants’ eyes and prevent visual fatigue, visual stimuli were rendered in narrow white outlines on black background. A short behavioral test given before the experiment confirmed that all participants could easily discriminate within novel pairs stimuli similar to those used in the experiment. Thus, it is unlikely that during the experiment participants made any substantial number of errors caused by perceptual difficulties.

**Figure 1.**
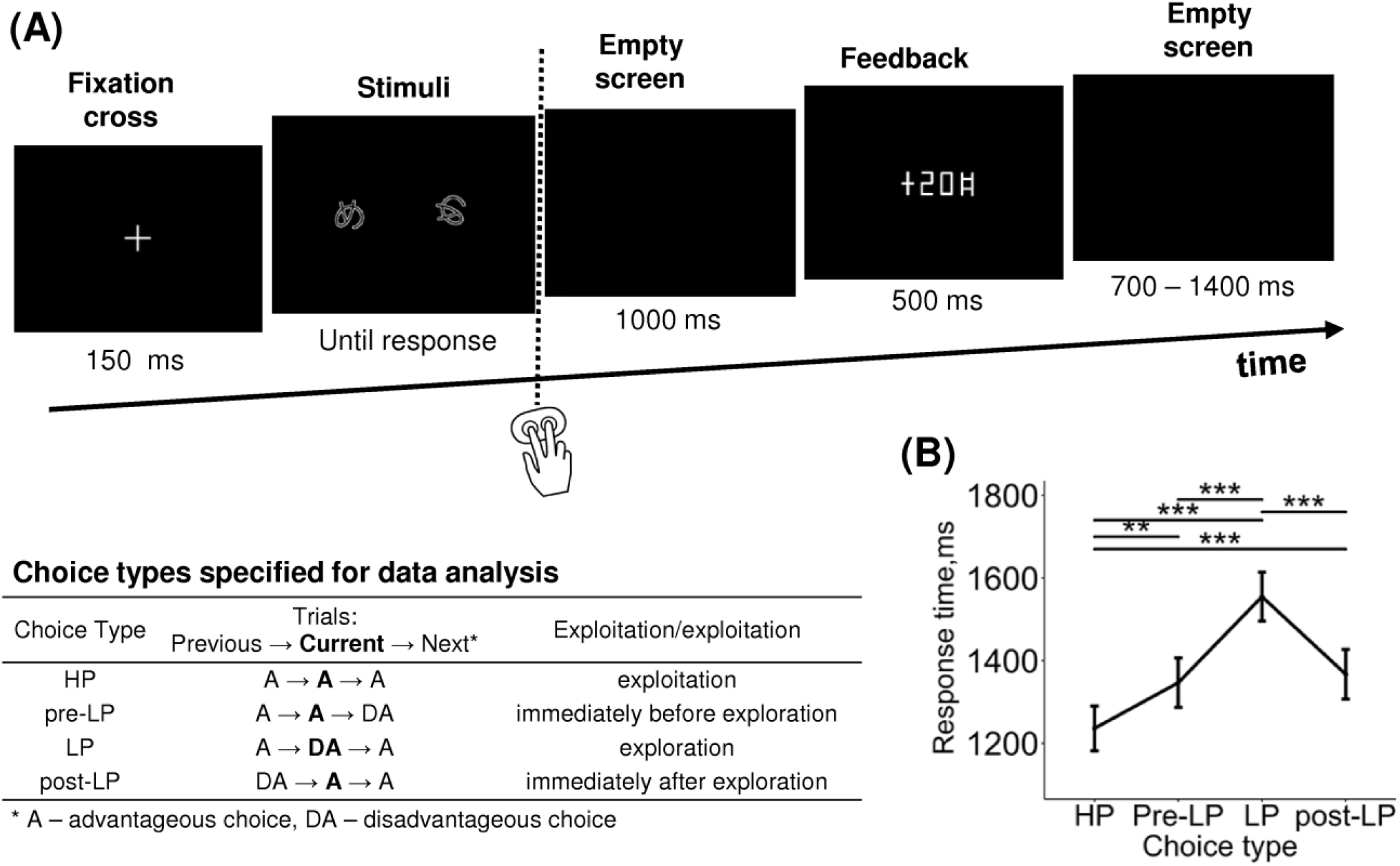
Probabilistic two-alternative gambling task: experimental design and behavioral statistics. **(A)** Experimental procedure. After a briefly presented fixation cross, a pair of visual stimuli was displayed on the screen. Participants learned by trial- and-error to select (by pressing one of the two buttons) the more advantageous of the two options, i.e., the one bringing monetary gains more often than the disadvantageous option, using the probabilistic feedback. After each choice, the number of points earned was displayed with a 1000-ms delay. Duration of each event is indicated under respective screens. The trials were organized into six blocks with different stimuli and reward structures between blocks. In each block, we took into analysis only ‘after learning’ periods, during which participants exhibited a stable behavioral preference for the advantageous stimulus. The table explains the further division of advantageous and disadvantageous choices into four choice types used in further analyses: high-payoff (HP) – advantageous/exploitative choices committed in a row of similar advantageous choices, low-payoff (LP) – rare disadvantageous/explorative choices, pre-LP and post LP – advantageous choices that were committed immediately before and immediately after an explorative LP choice. See text for details. **(B)** Response time as a function of choice type. Points and error bars on graphs represent M ± SEM across single trials in all subjects. ** – p<0.01, *** – p<0.001 (Tukey HSD).

Participants were instructed to make two-alternative choices between two stimuli presented on the screen simultaneously (a new pair was used in each block). Within each block, one of the stimuli was associated with greater probability of better outcomes (monetary gains on 70% of trials in blocks 1-5, and 60% of trials in block 6, with the rest of the trials leading to losses) resulting in higher average payoff – we will refer to them as ‘advantageous stimuli’. The other stimulus was associated with greater probability of worse outcomes (monetary gains on 30% of trials in blocks 1-5, and 40% of trials in block 6, with the rest of the trials leading to losses) resulting in lower average payoff; we will refer to them as ‘disadvantageous stimuli’. Outcome probabilities were constant within the blocks.

The sequences of gains and losses assigned to the stimuli throughout the experiment were generated quasirandomly in an interleaved manner that prevented multiple successive repetitions of the same outcome.

Prior to the experiment, we informed participants that one of the stimuli was more advantageous than the other. However, no further information regarding the stimuli was provided: participants were supposed to acquire knowledge about contingencies between their choices and outcomes by trial and error learning.

Each trial began with presentation of the fixation cross for 150 ms (Figure 1A), and then stimuli were presented on the screen continuously until a button was pressed by the participant. We did not restrict the response time aiming to reduce the number of impulsive decisions.

Participants’ behavioral responses were recorded with the use of a handheld MEG-compatible fiber optic button response pad (CurrentDesigns, Philadelphia, PA, USA). In order to indicate the choice of one of the two stimuli presented on the screen, participants pressed one of the two buttons using their index and middle fingers. Immediately following the button press, the screen was cleared.

Visual feedback was presented for 500 ms on each trial, with a delay of 1000 ms after the behavioral response; feedback informed participants about the number of points gained or lost on the current trial. The points were accumulated throughout the experiment. At the end of each block, current cumulative score was demonstrated. During the intertrial interval, the screen was empty (black). The intertrial interval varied from 700 to 1400 ms in a quasi-random order (flat distribution). By keeping the intertrial interval relatively short, we minimized the duration of the experiment in order to avoid fatigue and boredom in the participants.

The number of points associated with losses and gains was different across the blocks. In the first five blocks, which involved 70% and 30% reinforcement probabilities, we used five reinforcement schemes with the following magnitudes of losses and gains: (I) -20/+20, (II) 0/+20, (III) -20/0, (IV) +20/+50, (V) -50/-20, correspondingly. In the sixth block, which involved a lesser difference in reinforcement probabilities (60% and 40%), the reinforcement scheme was (VI) -20/+20. We used non-identical reinforcement schemes in successive blocks in order to make the blocks appear different for participants and thereby to make the participants learn in each block. We counterbalanced the order of reinforcement schemes across participants and used three sequences: I-II-III-IV-V-VI, I-IV-III-V-II-VI and I-IV-III-II-V-VI. We used only these three particular sequences for two reasons. First, in the initial block, we always used only mixed contingencies involving symmetrical magnitude of losses and gains (scheme I with -20/+20 points) in order to let the participants learn the effective choice strategy in the easiest way by comparing their cumulative gain with zero. Second, in the next block, we used only positive reinforcement schemes involving no absolute losses (+20/+50 or 0/+20 points). Thus, we ensured that participants earned a sufficient number of points during the first blocks of the experiment, providing for a non-negative total score throughout the experiment. This was needed in order to avoid frustration and loss of interest in participants. We converted the total score into rubles at a 1:1 ratio. On average, participants received 420 ± 250 rubles (mean ± SD).

Each of six experimental blocks comprised 40 trials with the overall duration of approximately 5 minutes per block. A short rest for approximately 1 minute (or longer if participants requested) was introduced between blocks. In total, the experiment lasted approximately 35 minutes.

The experiment was carried out using the Presentation 14.4 software (Neurobehavioral systems, Inc., Albany, CA, USA).

### 2.3 MEG data acquisition

MEG was recorded in a magnetically shielded room (AK3b, Vacuumschmelze GmbH, Hanau, Germany). We used the dc-SQUID Neuromag VectorView system (Elekta-Neuromag, Helsinki, Finland) with 204 planar gradientometers and 102 magnetometers. Sampling rate was 1000 Hz, and the passband was 0.03–330 Hz.

We measured participants’ head shapes by means of a 3Space Isotrack II System (Fastrak Polhemus, Colchester, VA, USA); we digitized three anatomical landmark points (nasion, and left and right preauricular points) as well as approximately 60 randomly distributed points on the scalp. Participants’ head position during MEG recording was continuously monitored by means of four head position indicator coils.

We used two pairs of electrodes placed above and below the left eye, and at the outer canthi of both in order to record vertical and horizontal electrooculogram, correspondingly. We also recorded bipolar electromyogram from the right neck muscles in order to identify muscular artifacts. For all recorded signals, sampling rate was 1000 Hz.

### 2.4 MEG data preprocessing

We applied the temporal signal space separation method (tSSS) (Taulu et al., 2005) to the raw data using MaxFilter (Elekta Neuromag software). The tSSS method suppresses magnetic interference coming from distant sources regarding the sensor array and thus can be used to remove biological and environmental magnetic artifacts. Data were converted to a standard head position (x = 0 mm; y = 0 mm; z = 45 mm). Static bad channels were detected and excluded from further processing steps.

For further offline analysis, MNE-Python software was used (Gramfort et al., 2013). We removed cardiac artifacts and artifacts related to eye movements from continuous data using the ICA method implemented in MNE-Python software.

In order to reduce computational cost and time required for time-frequency analysis, MEG data were resampled offline to 300 readings per second; this was safely above the Nyquist frequency since we were interested in MEG oscillations at much lower frequencies not exceeding 30 Hz.

We extracted response-locked epochs from -1750 to 2750 ms relative to the button press.

Correction of myogenic activity was performed using custom Python scripts with the use of MNE-Python functions. Specifically, we excluded contaminated epochs by thresholding the mean absolute signal values filtered above 70 Hz from each channel below 7 standard deviations of the across-channel average; such trials comprised 5.4 % of the experimental data.

### 2.5 Trial selection

First, we distinguished ‘advantageous’ choices (choosing the option with greater probability of gains) and ‘disadvantageous’ choices (choosing the option with lower probability of gains).

In the current study, we primarily sought to address directed exploration, i.e. voluntary goal-directed commission of disadvantageous choices purposefully aimed at gaining information (Wilson et al., 2014; Wilson et al., 2021). We assumed that directed exploration can be ascertained only if a subject has acquired a utility model of the task (i.e. has gained experience leading to effective usage of task contingencies), and acquisition of such a model would be expressed as a bias towards selecting the more advantageous option. Otherwise, exploration would likely be random rather than directed.

Thus, within each block, we extracted only those trials that met the following learning criteria: (1) these trials went after four consecutive choices of the advantageous stimulus and until the end of the block (the probability of three advantageous choices immediately following one by chance is (½)^3^ = 1/8 = 12.5%); (2) the percentage of advantageous choices within the selected trials was above 65%. This threshold was chosen because in a sequence of 30 trials (i.e. approximately the number of trials taken into consideration within an experimental block following the first step), 65% differs from the random level (50%) at the margin of significance (one-tailed one-sample binomial test p=0.05).

A recent pupillometric study (Kozunova et al., 2022) revealed that advantageous choices that immediately preceded and immediately followed explorative choices significantly differed from the advantageous choices committed within the periods of continuous exploitation. This finding hints that the internal state related to exploration modulates brain activity on a scale longer than duration of one trial. In the current study, we used the same approach, and in addition to distinguishing between advantageous and disadvantageous choices, we elaborated the trial classification that accounted for transitions between choice types in successive trials. Thus, for further analyses, we used the following four levels of Choice type factor (Figure 1; see also Supplementary Table S1): (1) the ‘high-payoff’ choice (‘HP’) – the advantageous choice that was preceded and followed by advantageous choices (such sequences of uninterrupted advantageous choices were considered as a stable ‘exploitative’ preference for the advantageous stimulus); (2) the trial preceding the ‘low-payoff’ choice (‘pre-LP’) – the advantageous choice preceding a disadvantageous decision and following an advantageous one; (3) the ‘low-payoff’ choice (‘LP’) – the disadvantageous ‘explorative’ choice that was preceded and followed by advantageous choices; (4) the trial following the ‘low-payoff’ choice (‘post-LP’) – the advantageous choice that followed a disadvantageous decision and preceded an advantageous one.

In addition, we classified trials depending upon the feedback received on the current trial (Feedback factor, two levels: ‘loss’, i.e. the worse of the two outcomes, and ‘gain’ i.e. the better of the two outcomes). For an additional follow-up analysis, we similarly classified trials depending on the feedback received during the previous trial (Previous feedback factor, two levels: ‘previous loss’ and ‘previous gain’).

Several participants were excluded from the analyses because of the lack of advantageous choices satisfying the learning criterion (2 participants) and due to failures to observe the behavioral procedure during recording (2 participants). Additionally, in order to keep the design more balanced, we excluded participants that had no trials in at least one of the eight conditions Choice type x Feedback (16 participants). Thus, statistical analyses reported below were done in 40 participants.

Trials with extreme response time (RT) values (<300 ms and >4000 ms) were excluded from all analyses; such trials comprised 1.4 % of the experimental data.

After rejection of epochs with extreme RTs and artifact-contaminated epochs, the number of epochs taken into analysis was 2908 in HP condition, 520 in pre-LP condition, 436 in LP condition, and 500 in post-LP condition.

We evaluated the dependence of RT upon choice type using the following LMM model, which included Choice type factor (four levels: ‘HP’, ‘pre-LP’, ‘LP’, ‘post-LP’), Previous feedback factor, i.e. outcome of the previous trial (two levels: ‘previous loss’ and ‘previous gain’) and their interactions as the fixed effects, and Subject as a random factor:

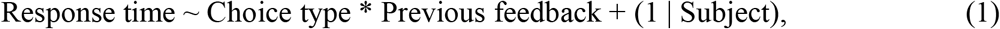

where Response time is log-transformed RT (time from stimulus onset to button press originally measured in milliseconds). We included the Previous feedback factor into the LMM model because it could affect the response time via interaction with Choice type in the same probability task (Kozunova et al., 2022).

### 2.6 Time-frequency analysis of the MEG data at the sensor level

We restricted our analyses of the β frequency range to 16-30 Hz (Engel and Fries, 2010; Kilavik et al., 2013). Frequencies below 16 Hz were not included into the analyses because the range of alpha-oscillations in MEG data may extend up to 15 rather than 12-13 Hz (Mierau et al., 2017).

We performed all the analyses using MNE Python open-source software (Gramfort et al., 2013) and custom scripts in Python. At sensor level, all time-frequency analyses were performed at single-trial level (unless stated otherwise), for each planar gradiometer independently. Discrete prolate spheroidal sequences (DPSS) multitaper analysis was applied to estimate spectra; this method is efficient for small numbers of trials (Thomson, 1982). This procedure increases signal to noise ratio at the cost of slightly diminished spectral resolution in a controlled manner. We used multitaper spectral analysis, implemented in the ‘mne.time_frequency.tfr_multitaper’ function (MNE-Python software, v. 0.21). The number of cycles was set as frequency divided by 2, thus the width of the sliding window was 500 ms at all frequencies, and it was shifted at steps of 25 ms. Time-bandwidth parameter was set to 4 (default value), resulting in frequency smoothing of 8 Hz.

Multitaper spectral analysis was applied at frequencies increasing in 2-Hz steps; we calculated β power within the 16 to 30 Hz range by summation of power data within the respective bands. After that, β power was log10-transformed and multiplied by 10 at each time point in each epoch and in each sensor. Log-transformation on single epochs has been demonstrated to render near-normally distributed EEG power across epochs and participants and thus to improve statistics performed on EEG power data (Smulders et al., 2018).

For baseline correction, we extracted baseline values from stimulus-locked epochs in the HP condition; we calculated β power in the time window from -350 to -50 ms relative to fixation cross onset using the same procedure as described above. Particularly, the MEG power data (log-transformed and multiplied by 10) were averaged across time points within the baseline window and across all trials belonging to the HP condition (for each participant in each sensor separately). Then, the baseline value was subtracted from β power values at all time points, for each participant in each sensor in each epoch independently. Thus, we obtained β power values relative to baseline expressed in decibels (dB).

We derived the baseline level from one of the conditions and used it to correct data within all conditions; the rationale was as follows. First, the intertrial intervals in the current experiment were rather short: if a traditional baselining procedure were used, carryover of the effects into intertrial intervals was likely to bias the analyses (cf. Kozunova et al., 2022). Second, tonic effects were reported in relation to the exploration-exploitation dilemma (Gilzenrat et al., 2010; Jepma et al., 2010; Jepma and Nieuwenhuis, 2011), and systematic tonic variations locked to explorative behavior could also bias a conventional pretrial baseline. Third, potentially, the decision to explore may be taken by participants during the intertrial interval rather than during the trial itself (since such a decision to explore per se does not require the participant to see the stimuli). Thus, the potential likelihood that a corresponding brain activity would emerge before the stimulus onset on explorative trials also precluded us from using the conventional pretrial baselining procedure. By using one baseline for all conditions, we were able to make unbiased comparisons between conditions. Such an approach is similar to a baseline-free approach, while it provides adjustment for spurious variability in the overall β power between subjects. We chose specifically the HP condition for baseline calculation because this condition represented persistent exploitation and it was probably the most stationary condition, as it involved sequential advantageous choices that were made in accordance with the participants’ internal utility model.

In order to reduce the dimensionality of the data and thus to increase statistical power under multiple comparisons, we combined sensors by adding time-frequency data within pairs of orthogonal planar gradientometers. Thus, at sensor level, we report power estimates for 102 such combined gradientometers; for brevity, we refer to them as sensors.

### 2.7 Statistical analysis

In order to estimate significant changes in the power of β oscillations over choice types and feedback valence, we applied linear mixed models (LMM) using the ‘lmer’ function available in the lme4 package for R (Bates et al., 2015). We checked the data for normality of the distribution of residuals with the normal quantile plot. To check the data for homoscedasticity we drew the residual plot.

We used linear mixed effects models (LMM) at a single-trial level rather than repeated measures ANOVA at the grand-average level for several reasons. The LMM method is robust to imbalanced designs – thus, missing data can be handled without listwise deletion of cases; moreover, different numbers of trials per condition are less problematic than in traditional ANOVA applied at the grand-average level (e.g., Kliegl et al., 2011). LMM is suitable for large numbers of repeated measurements per participant, thus making LMM especially appropriate to analyze data from individual trials: such a statistical approach allows accounting for intertrial variability, while standard averaging approaches preclude from using this informative data variability (Vossen et al., 2011; Tibon and Levy, 2015).

We used a data-driven approach towards selection of time windows and locations. Thus, the analysis was performed in two steps, as described below. At the first step, we used a LMM model aiming to identify salient effects with their time windows and clusters of sensors (with application of corrections for multiple observations). At the second step, we averaged β power within time windows and within significant sensors, and performed an in-depth data analysis, also using LMM statistics.

### 2.8 Defining time windows and clusters of sensors for further analyses

At the first step of our data-driven study, we used the following procedures on response-locked epochs at a single-trial level:

1. We averaged power of β oscillations over consecutive 200-ms windows centered on time points starting from -800 ms and up to 2400 ms relative to behavioral response onset (with feedback starting at 1000 ms), for each sensor independently. This procedure yielded 17 timeframes per epoch.
2. We applied a linear model with mixed effects (LMM) to single-trial data at each of 17 time points within each of 102 sensors independently. The model included Choice type factor (four levels: ‘HP’, ‘pre-LP’, ‘LP’, ‘post-LP’), Feedback factor (two levels: ‘loss’ and ‘gain’) and their interactions as fixed effects, and Subject as a random factor:

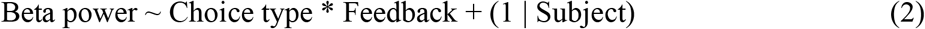
3. We applied correction for multiple comparisons using the false discovery rate (FDR) method (Benjamini and Yekutieli, 2001) to 1734 observations (17 time points x 102 sensors). At this stage, we plotted significant sensors (q=.05) on sequences of topographical maps, for each factor and their interaction.
4. Next, guided by our hypotheses and by visual inspection of the topographical plots with significant sensors indicated, we selected time windows, during which spatial distribution of significant sensors comprised distinct yet relatively stationary patterns.
5. For further analyses, we considered only those sensors that were persistently significant throughout all timeframes within the selected time windows. Additionally, in order to obtain compact and comprehensible clusters of significant sensors, we removed from analysis any significant sensors that had less than two significant neighbors.
6. In order to make sure that the effects to be analyzed represented either ERD or ERS, we retained for further analyses only those sensors that were significant against baseline for all conditions pooled (t-test, p <0.05, uncorrected). If within the given time window some sensors manifested ERD while other displayed ERS, we assigned two separate clusters of sensors correspondingly, thus separating two functionally distinct effects.

Additionally, for illustrative purposes, we rendered topographical plots representing grand-averaged spatial distributions of band power for consecutive 200-ms time windows (see Supplementary Figure S1).

In all illustrations of β power, when we considered choice types irrespective of feedback (i.e. for the two feedback valences pooled), we used the following two-step averaging procedure, for each subject, within each choice type, for each sensor separately. First, we averaged the epochs within each feedback valence, and then we averaged between the two feedback valences (in 1:1 ratio). Such a procedure was needed because the data matrix was not balanced: gains happened more often than losses after advantageous choices, while for disadvantageous choices losses predominated over gains. Thus, a simple flat averaging procedure across all trials within a particular choice type would lead to strong biases, while the two-step procedure allowed us to correct for the interdependence between Choice type and Feedback factors by equalizing the contribution of the Feedback factor. After that, the data were grand-averaged between participants.

### 2.9 Statistical analysis within the selected effects

After completing the first step that pinpointed plausible effects, we proceeded to the in-depth analysis of each effect separately, aiming to find out which particular pairwise contrasts created significant effects identified at the previous stage. At this step, for each selected effect separately, we collapsed MEG data across all timeframes within a corresponding time window and across sensors within the corresponding cluster, thus obtaining one value of β-band power per each trial.

For the analysis of effects preceding feedback onset, we applied to single-trial data the following LMM with the Choice type factor as the fixed effect, and Subject as a random factor:

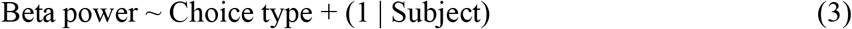

For effects following feedback onset, we applied to single-trial data the LMM that included Choice type factor, Feedback factor and their interactions as fixed effects, and Subject as a random factor:

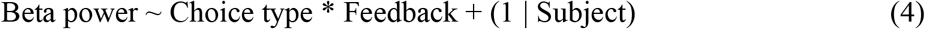

At this stage, we did not focus on the main effects and their interaction, which expectedly mirrored the findings obtained at the previous stage (see Supplementary Table S3). Instead, we aimed to find out which exactly statistical contrasts created the effects found at the previous stage. Thus, we complemented the basic LMM analysis with post-hoc pairwise comparisons using Tukey honestly significant difference (HSD) test (emmeans package version 1.6.0) (Lenth et al., 2020).

For illustrative purposes, we complemented these analyses by topographical plots of β-band power representing the most prominent pairwise contrasts between conditions, with FDR correction for 102 sensors (using the procedure similar to that described above). Additionally, we plotted timecourses of grand-averaged baseline-corrected β-band activity as well as event-related spectral perturbations for the β range and adjacent frequency ranges, for the data averaged over three most significant sensors within a particular cluster of sensors (see Supplementary Figure S2-S4).

In order to ascertain whether the power of β oscillations preceding the behavioral response reflects greater cognitive load during decision-making, we evaluated regressions at a single-trial level. We used the following LMM model:

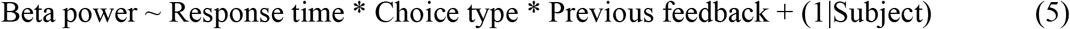

where Beta power is the power of β oscillations averaged over the time window within corresponding significant sensors (see above), and Response time is log-transformed time from stimulus onset to button press (originally measured in milliseconds; logarithmic transformation was needed because the distribution of response times deviated from normality).

Finally, we aimed to evaluate how the immediate history of losses and gains on the previous trial affected the brain response to the feedback on the current trial. We addressed the influence of the previous feedback because it was recently demonstrated that it affects behavioral and pupillometric measures on the next trial (Kozunova et al., 2022). Thus, for the time interval related to feedback processing, we evaluated joint effects of the feedback received on the current trial and the feedback received on the previous trial. This analysis was similar to the analyses described above, with the only exception that an additional factor Previous feedback corresponding to the outcome of the previous trial (two levels: ‘previous loss’ and ‘previous gain’) was included into the LMM model:

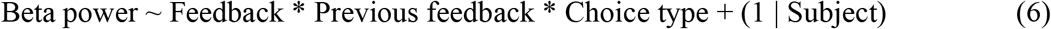

We evaluated statistical significance of pairwise differences using the post hoc Tukey honestly significant difference (HSD) test.

### 2.10 Source level analysis

Participants underwent MRI scanning with a 1.5T Philips Intera system (voxel size 1 × 1 × 1 mm, T1-weighted images). We reconstructed single-layer (inner skull) boundary-element models of cortical gray matter using individual structural MRIs, with a watershed segmentation algorithm (Ségonne et al., 2004) implemented in FreeSurfer 4.3 software (Martinos Center for Biomedical Imaging, Charlestown, MA, United States). To create individual anatomical surfaces, we used the ‘recon-all’ reconstruction algorithm implemented in FreeSurfer (Fischl et al., 2002) with default parameters. We co-registered head shapes into mesh using fiducial points and approximately 60 additional points on the scalp surface.

We used the ‘mne.setup_source_space’ function implemented in the MNE-Python open-source software (Gramfort et al., 2013) to create the source space, assuming that the sources are localized on the surface of the cortex. In order to compute the source space for the β-band power (16-30 Hz), we used ‘mne.minimum_norm.source_band_induced_power’ function (implemented in the MNE-Python open-source software), with Morlet wavelets (8 cycles per wavelet); the sliding window was shifted at steps of 25 ms. All epochs corresponding to each of the eight conditions (4 choice types x 2 feedback valences) were fed into this function within a single ‘mne.Epochs’ instance (implemented in the MNE-Python open-source software): thus, the resulting power estimates referred to the conditions rather than to single-trials.

We used a standardized low-resolution brain electromagnetic tomography (sLORETA) localization method (Pascual-Marqui, 2002). We used a grid with a spacing of 5 mm for dipole placement, which yielded 10,242 vertices per hemisphere. A noise-covariance matrix was computed for each participant from -350 ms to -50 ms relative to fixation cross onset in the HP condition. After that, β power was log10-transformed and multiplied by 10. Baseline correction was performed in the same way as at the sensor level: we used stimulus-locked epochs in the HP condition (from -350 to -50 ms relative to fixation cross onset) to estimate β-band power using the procedure described above. Then the β-band power data were log10-transformed and multiplied by 10 within this time window at each time point in each sensor in each epoch. After that, we calculated the baseline value by averaging data across 12 time points within the baseline window and across all trials within the HP condition (for each participant in each sensor separately). Then, we subtracted the baseline value from β-band power values at all time points, in each sensor in each epoch. Thus, the resulting β power relative to baseline was expressed in dB.

Pre-computed data for individual participants’ surface source estimates were morphed to a common reference space using the ‘mne.compute_source_morph’ function (implemented in the MNE-Python software). We used the ‘fsaverage’ template brain provided by FreeSurfer as a common reference space (Fischl et al., 1999).

We illustrated the power of β oscillations at source level in two ways. First, to illustrate spatial cortical localization of β-ERD and β-ERS, we rendered baseline-corrected power of β oscillations for all conditions collapsed. For this, we averaged the data between all 8 conditions, and then grand-averaged across participants, for each vertex separately. To render the data values on the inflated brain surface, we used the ‘mne.viz.plot_source_estimates’ function implemented in the MNE-Python software.

Second, we used the averaged data to illustrate spatial cortical localization of the pairwise contrasts. For contrasts between feedback valences the data were taken directly as obtained from the ‘mne.minimum_norm.source_band_induced_power’ function, while for contrasts between choice types (when considered irrespective of feedback valence), we averaged data between two feedback valences, within each choice type separately (for each participant within each vertex separately). After that, for each participant and within each vertex separately, we subtracted data between the two conditions being contrasted (e.g., LP vs. HP, or losses vs. gains within the LP choice type). Next, we grand-averaged the resulting differential data across participants. Finally, we plotted the grand-averaged data on the inflated brain surface, representing only significant vertices. For this purpose, we assessed significance of the difference in the averaged β-band power, for each vertex independently (pairwise t-test, p<0.05, uncorrected).

## 3 Results

### 3.1 Behavioral data

In the current study, we were interested in directed exploration. We argue that disadvantageous choices can be ascertained as directed exploration only on condition that participants had acquired an appropriate utility model of the task (otherwise, they probably made most of their choices randomly). Thus, only trials satisfying the learning criteria were included in the analyses (for behavioral statistics, see Supplementary Table S1).

Factor Choice type was significant for response time (RT) (LMM, F(_3,3519_) = 17.09, p<0.001) (Supplementary Table S2). RT was significantly longer for the LP condition compared with all the other conditions, and RTs for pre-LP and post-LP conditions were significantly longer compared with the HP condition (Tukey HSD, all p’s<0.01) (Figure 1B).

### 3.2 Data-Driven Approach to Choose Time Windows and Areas of Interest for the Analysis of Beta-Band Power Modulations

We expected that (1) brain processes related to decision-making will be different depending on the upcoming choice type (e.g., exploration vs. exploitation), and that (2) feedback valence will be processed differentially depending on the choice type that has been committed (contrasts of losses vs. gains will be different for explorative and exploitative trials).

In order to locate the expected effects in time and space, we combined a hypothesis-driven approach with a data-driven one. At the first step, we averaged response-locked β-band power data over consecutive 200-ms windows, in each sensor independently. We subjected these data to a single-trial statistical analysis using linear models with mixed effects (LMM), with Choice type, Feedback and their interaction included in the model as fixed effects. Significance was FDR-corrected for 102 sensors x 17 time windows. We visualized significant sensors for each factor and their interaction on topographical maps represented in Figure 2.

**Figure 2.**
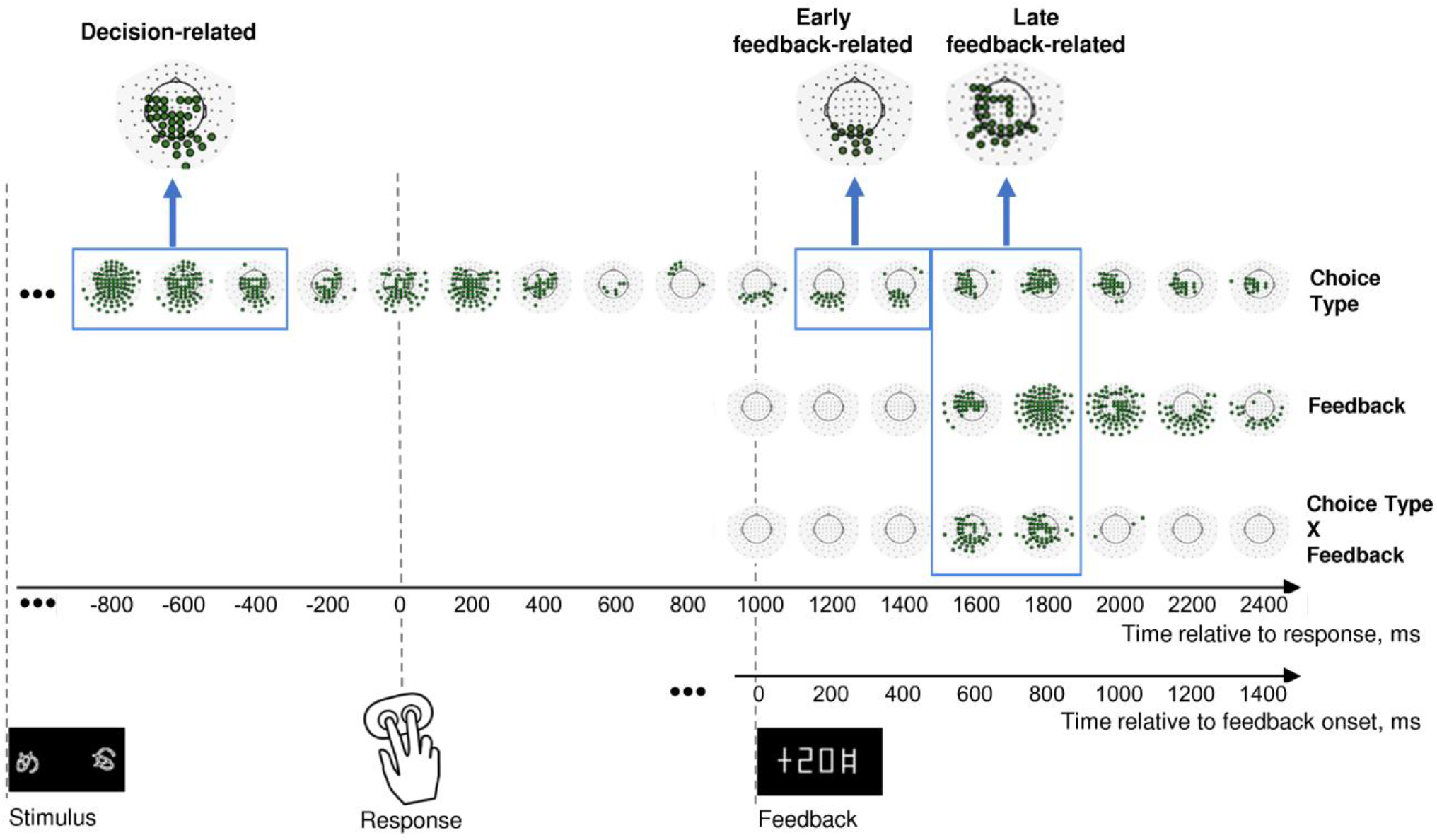
Effects of Choice type, Feedback, and their interaction on β power change in the consecutive 200 msec time intervals throughout an entire trial. Timelines at the bottom relate to response onset (zero point on the upper timeline) and to feedback onset (zero point on the lower timeline). The onsets of each event (stimulus presentation, choice selection, feedback) are marked by respective pictograms. Effect names are indicated at the right of the figure. The sequence of topographical maps corresponding to each effect demonstrates significant MEG sensors derived from LMM statistics (filled circles; p<0.05, FDR-corrected for 17 time windows x 102 sensors). Blue frames mark time windows used in further analyses of the effects (decision-related, early feedback-related, and late feedback-related), with inserts at the top of each rectangle representing clusters of sensors taken for analysis. See Methods for details.

In order to test our hypotheses, we focused on the three time intervals with corresponding sensor topography. A profound lasting effect of the Choice type factor was present before and during the behavioral response (Figure 2, upper row). Since we were interested in the brain processes related to decision-making leading to initiation of the behavioral response, we restricted the further analysis to the pre-response part of the effect within -900 – -300 ms relative to the response onset, thus aiming to avoid a possible overlap with effects related to stimulus processing and movement execution.

Feedback-related effects were not uniform, thus we selected two separate effects: (1) an early feedback-related effect of Choice type around the feedback onset at 100 – 500 ms after feedback onset (Figure 2, upper row); (2) a late feedback-related interaction of Choice type x Feedback at 500 – 900 ms after feedback onset (Figure 2, bottom row). The latter effect apparently evidenced differential feedback valence processing depending upon the choice type.

For data visualization on topographical maps, event-related spectral perturbations and timecourses, see Supplementary Figure S1-S4.

### 3.3 Decision-making effect

To further analyze the dependence of β power upon the choice type, we averaged β power across the cluster of sensors demonstrating the respective effect within the pre-response time window (Figure 2). Strong and highly significant suppression of β power relative to baseline was observed during the decision-making period (Figure 3A). A source-level analysis revealed the widespread β-ERD originating from the left frontal, left pericentral areas, as well as from the parietal, occipital (both lateral and medial aspects), and medial temporal areas (Figure 3E).

**Figure 3.**
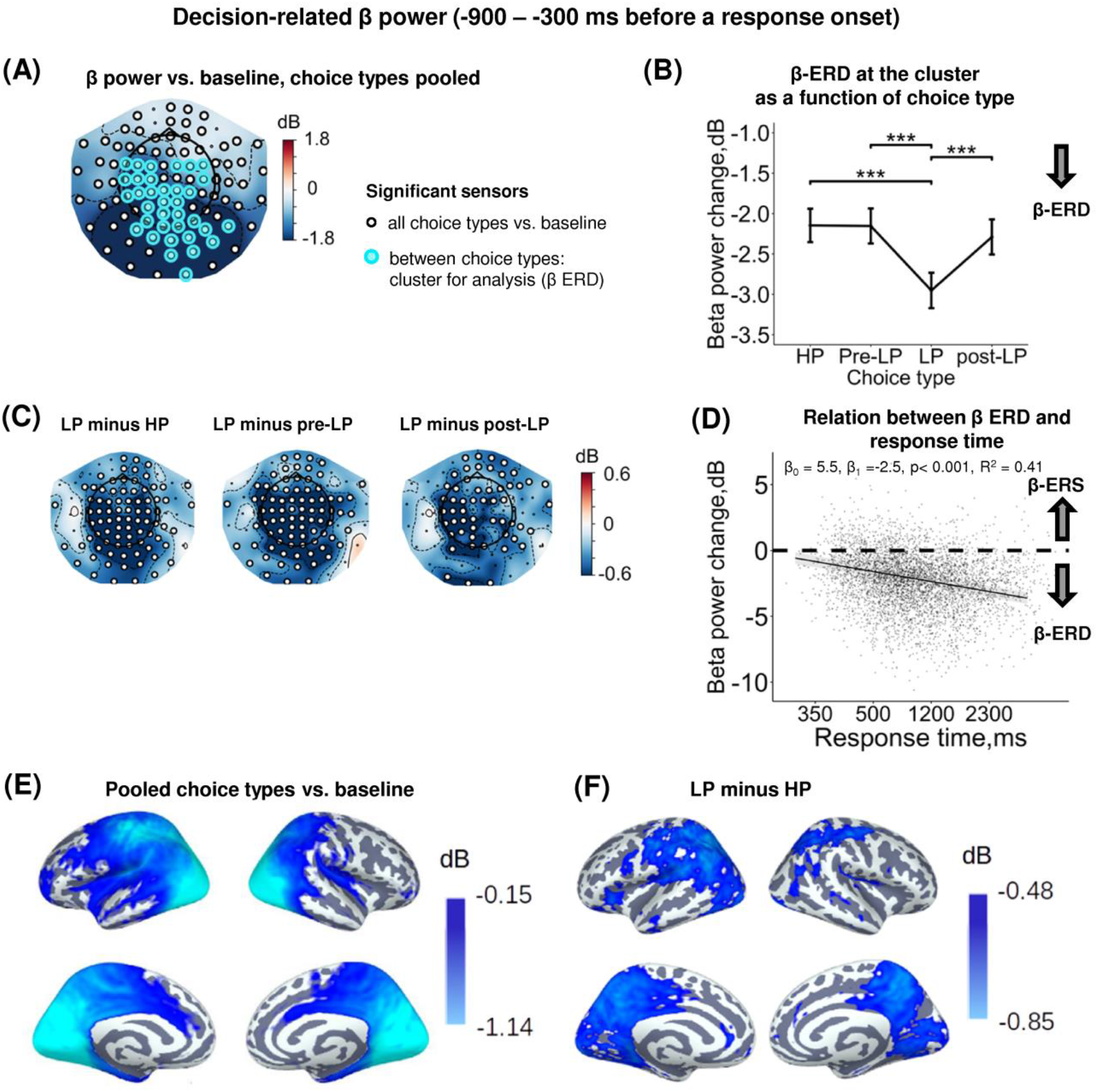
Enhanced decision-related β power suppression (β-ERD) distinguishes explorative LP choice from exploitative choice types. **(A)** Topographic map representing β-band power change relative to baseline averaged over the time interval of -900 – -300 ms before a response onset, all choice types pooled. Small black open circles mark sensors, in which β power is significantly different from baseline (t-test, p<0.05, uncorrected); sensors included in the cluster differentiating choice types according to LMM analysis (cf. Fig 3) are additionally marked by colored rings. Here and hereafter, blue and red colors denote β-ERD and β-ERS respectively. **(B)** Decision-related β-band power change averaged over the cluster of significant sensors (colored rings in panel A), as a function of Choice Type. The β power change for each choice type show averages across sensors per significant cluster marked by the colored rings in panel A. Points and error bars on graphs represent M ± SEM in single trials in all subjects. *** – p<0.001 (LMM, Tukey HSD test). **(C)** Topographic maps of the pairwise difference in β power change between the LP and other choice types at the sensor level. Significant sensors (LMM, Tukey HSD test, p<0.05, FDR-corrected) are indicated by small black open circles. **(D)** Linear regression of response time to decision-related β-band power change averaged across sensors of significant cluster (colored rings in panel A), all choice types pooled. The line represents linear regression; shaded areas depict 95% confidence interval. Dots on scatterplots correspond to β power change in individual trials in all participants. **(E and F)** Spatial distribution of cortical sources underlying decision-related β-ERD: (E) relative to baseline, all conditions pooled; and (F) increased β-ERD for LP as compared to HP choices. Only significant vertices are shown (t-test, p < 0.05, uncorrected). For event-related spectral perturbations and timecourses of β power of the LP vs. HP contrast, see Supplementary Figure S2.

At the sensor level, the effect of choice type (Supplementary Table S3) was due to deeper β-ERD in LP trials compared with all the other trial types (HP, pre-LP and post-LP) (Tukey HSD, all p’s<0.001) (Figure 3B, see also Figure 3C). At the source level, the significant difference between LP and HP conditions was localized to left pericentral, most of the parietal areas and large parts of the occipital cortex (both on lateral and medial aspects), as well as posterior cingulum (Figure 3F).

Thus, as we expected, decision-making regarding explorative choices involved deeper suppression of β oscillations compared with exploitative decisions.

In order to find out whether β suppression during decision-making is indeed related to greater cognitive load (which may explain delayed responding), we estimated regression between response time and β-band power. We observed a highly significant negative regression slope (LMM statistics, β_0_ = 4.82, β_1_ =-2.33, p< 0.001, R^2^ = 0.41) (Figure 3D). Negative slope indicates that the slower decision-making was the more β power was suppressed during decision-making, thus suggesting greater cognitive load during slower decision-making.

### 3.4 Early effect of choice type on the feedback processing

Starting around the time of the visual feedback onset and sustaining up to 500 ms, the effect of Choice type on β power was observed in the occipito-parietal cluster of sensors (Figure 2, Figure 4A and Figure 4D). For all choice types, early post-feedback β power in this cluster was suppressed relative to baseline (Figure 4 A, B). Note that the concurrent rise of β power (β-ERS) in the anterior sensors (Figure 4A) was not influenced by Choice type and started long before the feedback onset (Supplementary Figure S1), thus hardly representing the early feedback-related response. The effect of Choice type (Supplementary Table S3) was mainly driven by stronger β-ERD on LP trials compared with all the other trial types (Tukey HSD, all p’s <0.05) (Figure 4B, see also Figure 4C). Notably, the feedback valence (whether a participant gained or lost) did not influence β-ERD in any trial type (Supplementary Table S3). Source-level analysis of the LP vs. HP contrast revealed that the LP-related increase in the β-ERD mainly occupied cortical regions on the lateral and medial surface of the occipital and posterior part of parietal and occipital lobes (Figure 4E).

**Figure 4.**
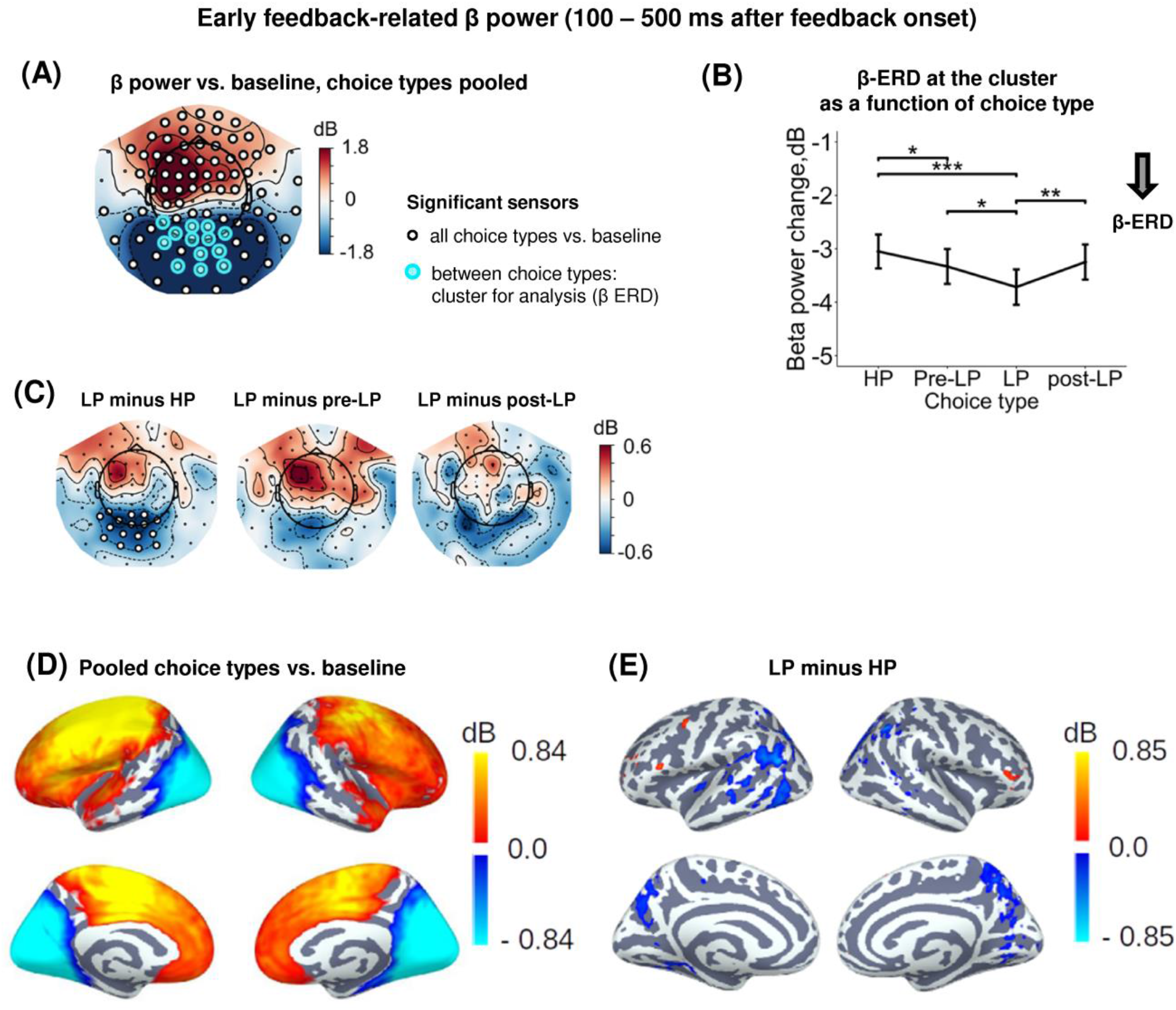
Enhanced early feedback-related β-ERD at posterior cortex distinguishes explorative LP choice from exploitative choice types. **(A)** Topographic map of feedback-related β-band power change relative to baseline averaged across the time interval 100 – 500 ms after feedback onset, all choice types pooled. Blue and red colors denote β-ERD and β-ERS respectively. Small black open circles mark sensors, in which feedback-related β power is significantly different from baseline (t-test, p<0.05, uncorrected); sensors included in the cluster differentiating feedback-related β power between choice types according to LMM analysis (cf. Figure 3) are additionally marked by colored rings. Note that β-ERD in the posterior cluster significantly varies between choice types, while β-ERS in the anterior sensors does not differentiate between choices. **(B)** Early feedback-related β-band power change averaged over the cluster of significant sensors (colored rings in panel A), as a function of Choice Type. Points and error bars on graphs represent M ± SEM across single trials in all subjects. * – p<0.05, ** – p<0.01, *** – p<0.001 (LMM, Tukey HSD test). **(C)** Topographic maps of the pairwise difference in early feedback-related β power change between the LP and other choice types at the sensor level. Significant sensors (LMM, Tukey HSD test, p<0.05, FDR-corrected) are indicated by small black open circles **(D and E)** Spatial distribution of cortical sources underlying feedback-related β power change: (E) relative to baseline, all choice types pooled; and (D) increased β-ERD for LP as compared to HP choices. Only significant vertices are shown (t-test, p < 0.05, uncorrected). For event-related spectral perturbations and timecourses of β power of the LP vs. HP contrast, see Supplementary Figure S3.

Thus, as we predicted, explorative choices boosted processing of the visual feedback signal, no matter whether their outcome was good or bad for a participant.

### 3.5 Late feedback-related effect

A significant Choice type x Feedback interaction distinguished the late feedback-related time interval (500 – 900 ms after feedback onset) (Figure 2). This implies that, as we expected, β power modulations by positive and negative feedback depended upon choice type. Yet, surprisingly, this dependency manifested itself relatively late throughout the course of feedback-related brain response. For all trial types pooled together, we observed a significant β-ERS in the anterior areas, while significant β-ERD persisted over the posterior regions (Figure 5A and D). Therefore, we proceeded with analyses in sensor subclusters manifesting significant ERD and ERS separately.

**Figure 5.**
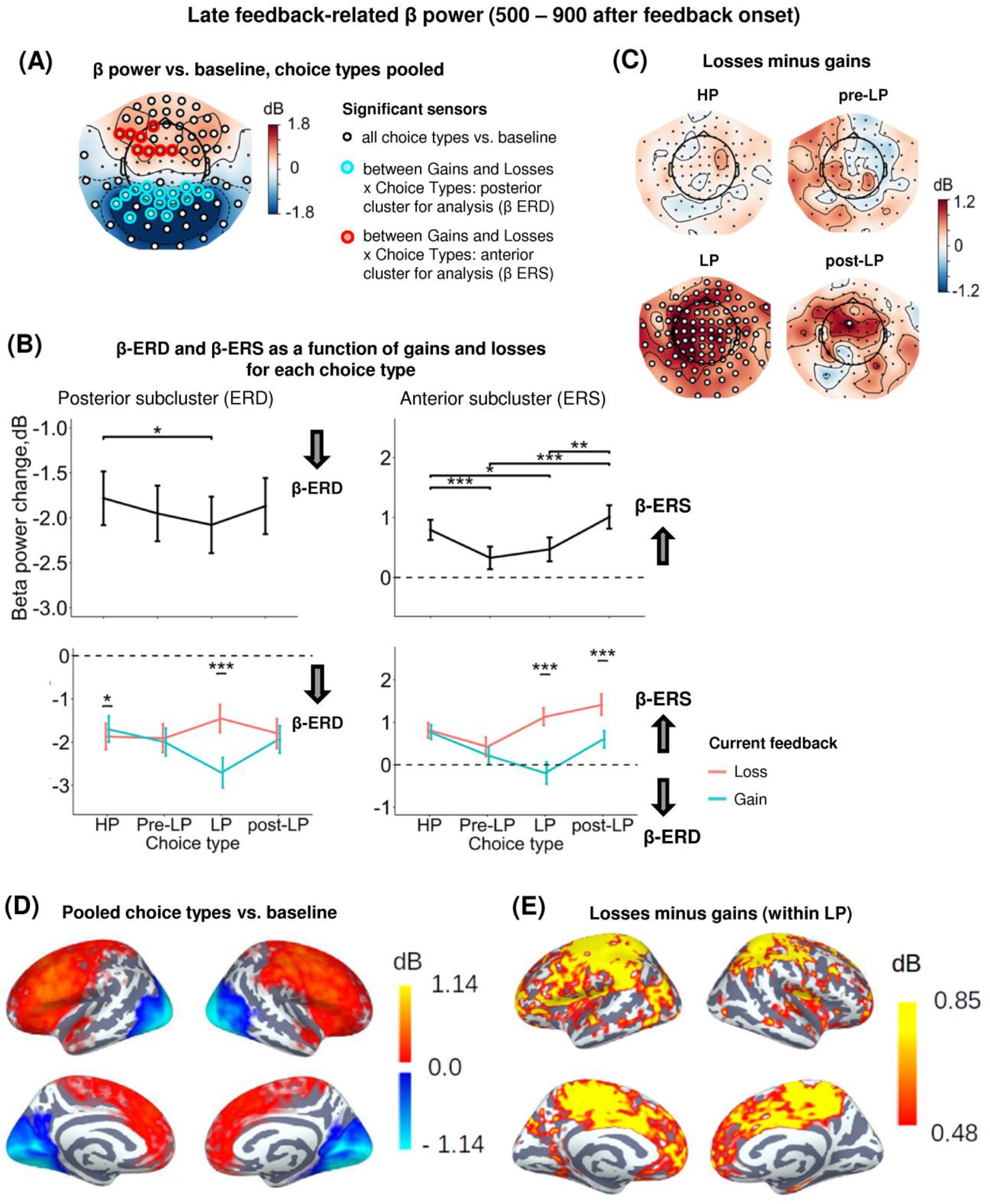
Heightened sensitivity to losses versus gains in late feedback-related anterior β-ERS and posterior β-ERD distinguishes explorative LP choices from exploitative choice types. **(A)** Topographic map of late feedback-related β-band power change relative to baseline averaged over the time interval 500 -900 ms after feedback onset, all choice types pooled. Small black open circles mark sensors, in which late feedback-related β power is significantly different from baseline (t-test, p<0.05, uncorrected); sensors included in the cluster differentiating feedback-related β power following gains and losses between choice types according to LMM analysis (cf. Figure 3) are additionally marked by colored rings. **(B)** Late feedback-related β-band power change averaged over the clusters of sensors (colored rings in panel A): anterior cluster displaying β-ERS and posterior cluster displaying β-ERD, as a function of Choice Type (upper row) and its interaction with gain or loss (lower row). Points and error bars on graphs represent M ± SEM across single trials in all subjects. * – p<0.05, *** – p<0.001 (LMM, Tukey HSD test). **(C)** Topographic maps representing contrasts between losses and gains for each choice type separately. Significant sensors (LMM, Tukey HSD test, p<0.05, FDR-corrected) are indicated by small black open circles. **(D and E)** Cortical distribution of sources demonstrating β power change: (D) relative to baseline, all conditions pooled, and (E) the contrast between losses and gains within the LP choices. Only significant vertices are shown (t-test, p < 0.05, uncorrected). For event-related spectral perturbations and timecourses of β power of the losses vs. wins contrast under LP condition, see Supplementary Figure S4.

#### 3.5.1 Posterior subcluster

Similarly to the early post-feedback interval, a significant β-ERD relative to baseline was observed in the posterior sensors (Figure 5A, Figure 5D, Supplementary Table S3). For the posterior subcluster, LMM analysis confirmed the significance of Choice type, Feedback, and Choice type x Feedback interaction (Figure 5A, Supplementary Table S3). The posterior β-ERD was deeper for LP trails compared to HP trials (Figure 5B), i.e., it manifested the same, although less pronounced, LP-related bias as we observed in the early post-feedback interval. Choice type x Feedback interaction was mainly explained by significantly greater posterior β-ERD after positive compared with negative feedback on LP trials (Tukey HSD, p < 0.001), but not for the other choice types (Figure 5B, C). On the contrary, on exploitative HP trials sensitivity of posterior β-ERD to positive and negative feedback was opposite -with β-ERD being slightly yet significantly larger after losses compared with gains (Tukey HSD, p = 0.03) (Figure 5B).

Thus, selectively increased posterior β-ERD indexed greater neural activation after losses for serial exploitative HP choices and after wins in the explorative LP choices. Given the higher probability of wins after HP choices and losses after LP choices, these findings suggested that the activation strength depended on the outcome expectancy, with greater activation accompanying late processing of the less probable outcome.

#### 3.5.2 Anterior subcluster

Sensors of the anterior subcluster exhibited significant β-ERS (Figure 5A, Figure 5D, Supplementary Table S3). Factors Choice type, Feedback and interaction Choice type x Feedback were all highly significant (Supplementary Table S3). The effect of the Choice type can be explained by the fact that anterior β-ERS was significantly greater for HP trials compared with pre-LP and LP trials (Figure 5B). Since a choice of the advantageous stimulus has been rewarded more frequently than its risky alternative, this finding seemed to be in line with the role of the reward in anterior β-ERS induction for the specific, possibly, advantageous choices.

However, examination of Choice type x Feedback interaction by splitting the data by feedback valence (Figure 5B, see also Figure 5C) ruled out this explanation. Negative and positive feedback in the HP trials equally contributed to the anterior β-ERS, while in explorative LP trials β-ERS was driven exclusively by negative feedback (Figure 5B, see also Figure 5C and Figure 5E). Note that in the LP trials, positive feedback was even followed by β power suppression relatively to baseline – instead of its increase (Figure 5B).

In addition, trials immediately preceding explorative LP choices (pre-LP), despite their superficial similarity with the advantageous HP choices, showed a significantly reduced ERS to feedback (Tukey HSD, p < 0.001), regardless of the feedback valence (Figure 5B). Yet, the same advantageous choice in the post-LP trials, similarly to the LP trials themselves, induced greater anterior β-ERS to losses compared with gains (Tukey HSD, p < 0.001, Figure 5B).

The high sensitivity of anterior β-ERS to punishment in the explorative choices (and to a lesser degree in post-explorative choices) strongly suggests that this specific form of oscillations was not driven by the valence of the feedback signal itself, but rather depended on its context, which, in our task, varies between highly expected (serial HP trials) and that involving conflict with the accumulated reward history (LP trials).

These findings on β power sensitivity to feedback valence on explorative trials agree with the predictions made from the ‘predictive coding’ account of anterior β-ERS outlined in the Introduction. However, none of the predictions can account for the finding that increased sensitivity of anterior β-ERS to punishment would sustain in the post-LP advantageous choices when a participant came back to the non-conflicting exploitative strategy. This finding implied that in addition to the accumulated history of trials and errors, there could be a short-term impact of the previous trial outcome on the anterior β-ERS. We explored this possibility in the following section.

### 3.6 The impact of the feedback presented on the previous trial

In order to evaluate the impact of the previous feedback (i.e., that was presented on the previous trial) on the late feedback-related effect evoked by losses and gains on the current trial, we ran LMM analysis with Previous feedback fixed factor additionally included into the model (Figure 6). Specifically, we were interested only in ERS effects (i.e. the anterior subcluster) and two choice types – HP and LP.

**Figure 6.**
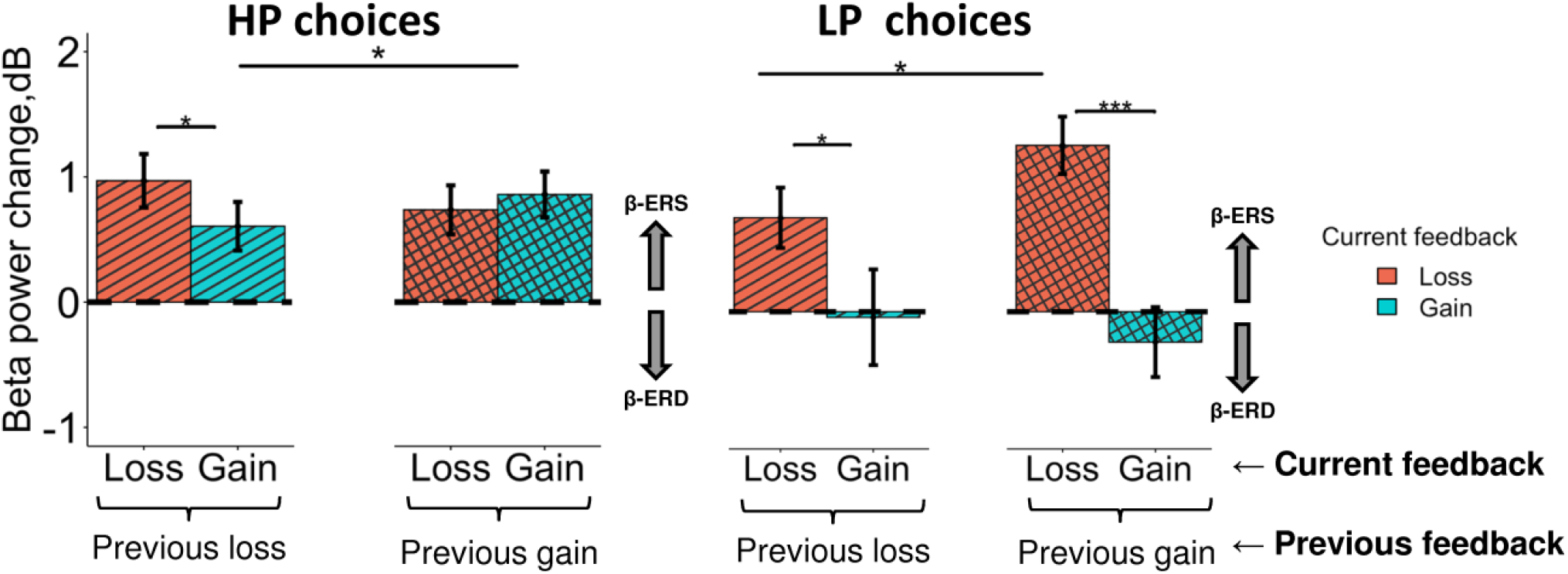
Different impact of the previous choice outcome on anterior β-ERS induced by losses and gains in the LP and HP choices. Late β band power change (500 – 900 ms after feedback onset) averaged across the anterior cluster (see Figure 5A) are shown for LP and HP choices as a function of two factors: a current feedback valence (‘loss’ and ‘gain’, red and blue rectangles respectively) and a previous feedback valence (‘previous loss’ or ‘previous gain’ indicated by horizontal brackets). Boxes and error bars on graphs represent M ± SEM across single trials in all subjects. Asterisks denote Tukey HSD p-values: *p<0.05, ***p<0.001.

Triple interaction Choice type x Previous feedback x Feedback was significant (F_1, 3304_ = 6.0, p = 0.015). For exploitative HP choices (Figure 6, left panel), β-ERS induced by a current gain was slightly but significantly enhanced if the previous similarly advantageous choice was expectedly rewarded rather than punished (Tukey HSD, p = 0.02), whereas β-ERS induced by a current loss remained unaffected by the previous outcome.

On the contrary, for the explorative LP choices, the outcome of the previous advantageous choice significantly affected β-ERS in the case of a current loss (Tukey HSD, p = 0.04) but not a current gain (Tukey HSD, p >> 0.05) (Figure 6, right panel). It should be stressed that gains for LP choices produced no β-ERS at all.

A strong β-ERS bound to anticipated monetary losses ensuing from objectively disadvantageous choices compared with gains, was highly significant if the previous advantageous choice was rightfully rewarded (Tukey HSD, p < 0.001), while after punishment it was at the margin of significance (Tukey HSD, p = 0.04).

In other words, for HP and LP choices similarly, anterior β-ERS was further enhanced if not only the outcome of the current choice but also the outcome of the previous one matched the prediction/expectation derived from the utility model acquired from the previous experience.

## 4 Discussion

Using a simple two-choice probabilistic reinforcement learning task, which was learnt by participants through trial and error, we investigated β-band power modulations related to rare disadvantageous choices conflicting with the acquired inner utility model.

The study brought two main findings. First, decision making leading to explorative choices took more time and evidenced greater large-scale suppression of β oscillations than its advantageous alternative. Crucially, the magnitude of a large-scale β suppression, when a subject formed his/her decision reliably, predicted decision costs (with response time taken as a measurable proxy of this internal variable) on a single trial basis. Given multiple evidence associating the strength of β suppression with greater cognitive and attentional efforts (Scharinger et al., 2017; Pavlova et al., 2019; Tafuro et al., 2019), this finding strongly suggests that such a choice requires additional resources to overcome the internal utility model favoring the advantageous alternative and strongly supports the hypothesis of its deliberately explorative nature (Ellerby and Tunney, 2017; Kozunova et al., 2022).

Second, the most intriguing finding concerns β oscillations induced by feedback received after deliberately explorative choices. The main difference between directed explorative and exploitative choices was the impact of their negative versus positive outcome on β oscillations arising during post-evaluation processes (Figure 5). When a choice was made within a sequence of similarly advantageous choices (HP choice), both monetary gains and losses produced a late synchronization of frontal β oscillations. These feedback-related β oscillations were reduced to a baseline if a participant was going to switch from such an advantageous strategy toward the explorative one (pre-LP choice). Remarkably, only losses – but not gains – were accompanied by reliable frontal β synchronization after committing directed exploration (LP choice).

Therefore, in our task, there was no consistent relationship between the presence or absence of frontal β oscillations and the reward or punishment for a committed choice. Therefore, merely maintaining the ‘reward-induced’ state, as the ‘reward’ hypothesis implies (Marco-Pallares et al., 2015), is not sufficient to generate enhanced β power. Although the context-dependency of punishment-related frontal β was previously mentioned by some authors (e.g., Yaple et al. (2018)), this conclusion was derived from the findings obtained in different research paradigms. Here, for the first time, we directly contrasted frontal β resulting from the outcomes of two choice types committed by the same participants within the same task on the trial-by-trial basis. The solely punishment-related β synchronization observed for explorative choices cannot be explained or predicted by previously suggested contextual factors such as successful punishment avoidance (Hamel et al., 2018), task difficulty (Billeke et al., 2020), emotional salience of the feedback signal (Leicht et al., 2013), reward expectancy (Cohen et al., 2007; Donamayor et al., 2012; HajiHosseini et al., 2012; Mas-Herrero et al., 2015; Ramakrishnan et al., 2017). It is possible that the bias of the literature toward a role of β oscillations in positive feedback processing derives in part from experimental biases, e.g., nonequivalence of the relevance of positive and negative outcomes for the future behavior (as in HajiHosseini and Holroyd (2015)) and/or a complete lack of response-feedback contingency, i.e., a random structure of rewards and punishments for either choice (HajiHosseini et al., 2012; Mas-Herrero et al., 2015).

Being contradictory to the ‘pure reward’ hypothesis, our results are instead consistent with the role of β oscillations in the brain state resulting from the matching processes between the outcome of a current choice and accumulated experience of gains and losses, i.e., the inner utility model.

Both positive and negative feedback signals used in our task could induce the increase in β power with a latency of five to nine hundred milliseconds. However, late β ERS clearly depended on the congruency between the feedback received and the prediction based on the previously acquired utility model. Specifically, when a participant’s choice complies with the inner utility model (i.e., if the advantageous stimulus with the higher objective reward probability was repeatedly chosen), late frontal β increases regardless of the actual feedback that a participant has received. Given the probabilistic nature of the model, the precision of its prediction is not very narrow (see e.g., Yon and Frith (2021)), and a single loss resulting from the objectively advantageous choice does not lead to a tangible mismatch. By contrast, explorative choices committed against the predominant value-based response tendency trigger frontal β synchronization only after a monetary loss, which provides strong evidence for validity of the inner utility model. In this case, a reward rather than punishment for a preceding advantageous choice, being also congruent with an inner model’s prediction, increases β-ERS induced by a current loss (Figure 6). In other words, for both exploitative and explorative choices, maximal frontal β-ERS accompanies those feedback signals that reliably confirm the validity of the existing value-based model – even if the current choice was driven by its explorative alternative. This type of β oscillations thus appears as a categorical information and not as a scalar measure of discrepancy between outcome and expectation.

The relatively long latency (500-900 ms) of the feedback-related frontal β-ERS places strong constraints on its potential functional significance. It is unlikely that the differential response of β-ERS to reward and punishment in our task is causally involved in the feedback valence evaluation, as it occurs substantially after the ‘feedback processing’ time window, which is usually between 200 and 500 ms relative to feedback onset (Cohen et al., 2007; Marco-Pallares et al., 2008); but see also Yaple et al. (2018) and Alicart et al. (2020), see also Leicht et al. (2013) for a later latency of differential effect of feedback valence on frontal β under specific task requirements).

We offer here that there is a more sophisticated account of feedback-related frontal β, which is focused on the post-evaluative memory-based replay of ‘cognitive set’ (Engel and Fries, 2010) or rule representation (Buschman et al., 2012; Brincat and Miller, 2015; Brincat and Miller, 2016) with its possible role in the strengthening of the re-activated cortical representation. In line with this idea, the cortical topography of the late frontal anterior β-ERS (Figure 5) reveals involvement of a widely distributed set of cortical areas that are mainly located on the dorsal and medial surface of the frontal cortex including anterior and dorsal cingulum, supplementary motor area, inferior frontal gyrus, anterior insula and ventromedial frontal cortex. All these structures constitute cortical nodes of the cingulate-basal ganglia-limbic circle, strongly implicated in the stabilization of neural representations for selected behavioral rule through reverberating activity in the β-range (Leventhal et al., 2012; Brincat and Miller, 2016).

Our proposal here is closely related to this idea, yet places it in a context of exploration-exploitation dilemma. We suggest that during outcome evaluation of a deliberately explorative choice, a representation of the previously rewarded choice strategy (utility model) is maintained in the memory buffer alongside with that for the voluntary decision, which shifts current choice priorities in favor of information seeking, i. e. testing the alternative model, or ‘task set’ (Domenech et al., 2020; Koechlin, 2020). Such competing representations are thought to be re-activated whenever the current task induces conflict in processing preceding response selection, but also in feedback evaluation (Botvinick et al., 2004). The evaluation process lends the greatest weight to the external feedback that is most reliable, i.e., matches the previously accumulated experience (see e.g., Yon and Frith (2021)), aka the inner utility model in our experimental settings. The further stabilization of the winning utility model is reflected in the punishment-related β oscillations for alternative explorative choice, and it would compromise future repetitions of the explorative behavioral program in the unchanging environment. A rare gain for such choices does not provide sufficient evidence to strengthen either model; therefore, it does not induce β synchronization. This leaves a window of opportunity for ‘set shifting’, i.e., for adjustment of value-based model in the future, which is vital in case environment changes. We predict that using the same task in the ‘reversal learning’ context will reveal selectively increased reward-related β oscillations for a number of explorative choices made after the reversal of previously acquired stimulus–response associations.

To conclude, our results suggest that emergence of late punishment-related β-ERS for deliberately explorative choices reflects a replay of a value-based model representation that possesses greater predictive capacity in an unchanging environment and will be strengthened for future use.

## 6 Author Contributions

G. L. K. and T.A.S. contributed to conception and design of the study. G. L. K. and A.O.P. conducted the experiments. B.V.C., K.I.P., V.D.T., and A.S.M. analyzed the results and prepared illustrations. B.V.C. and T.A.S. wrote the first draft of the manuscript. B.V.C., K.I.P., V.D.T., A.S.M., A.O.P., G.L.K., T.A.S. wrote sections of the manuscript. All authors reviewed the manuscript.

## 8 Acknowledgments

We are grateful to Ksenia E. Sayfulina for developing scripts for statistical analysis, and to Ksenia E. Sayfulina and Anastasia Yu. Nikolaeva for invaluable help with data collection.

This manuscript has been released as a pre-print at bioRxiv.

## 9 Data Availability Statement

The datasets generated and/or analyzed during the current study are available from https://figshare.com/projects/Losses_resulting_from_deliberate_exploration_trigger_beta_oscillations_in_frontal_cortex/149792

## 10 Code Availability Statement

The code (custom scripts) generated during the current study are available from the corresponding author on reasonable request.

