## Supplementary for "Losses resulting from deliberate exploration trigger beta oscillations in frontal cortex"

***Supplementary Material***

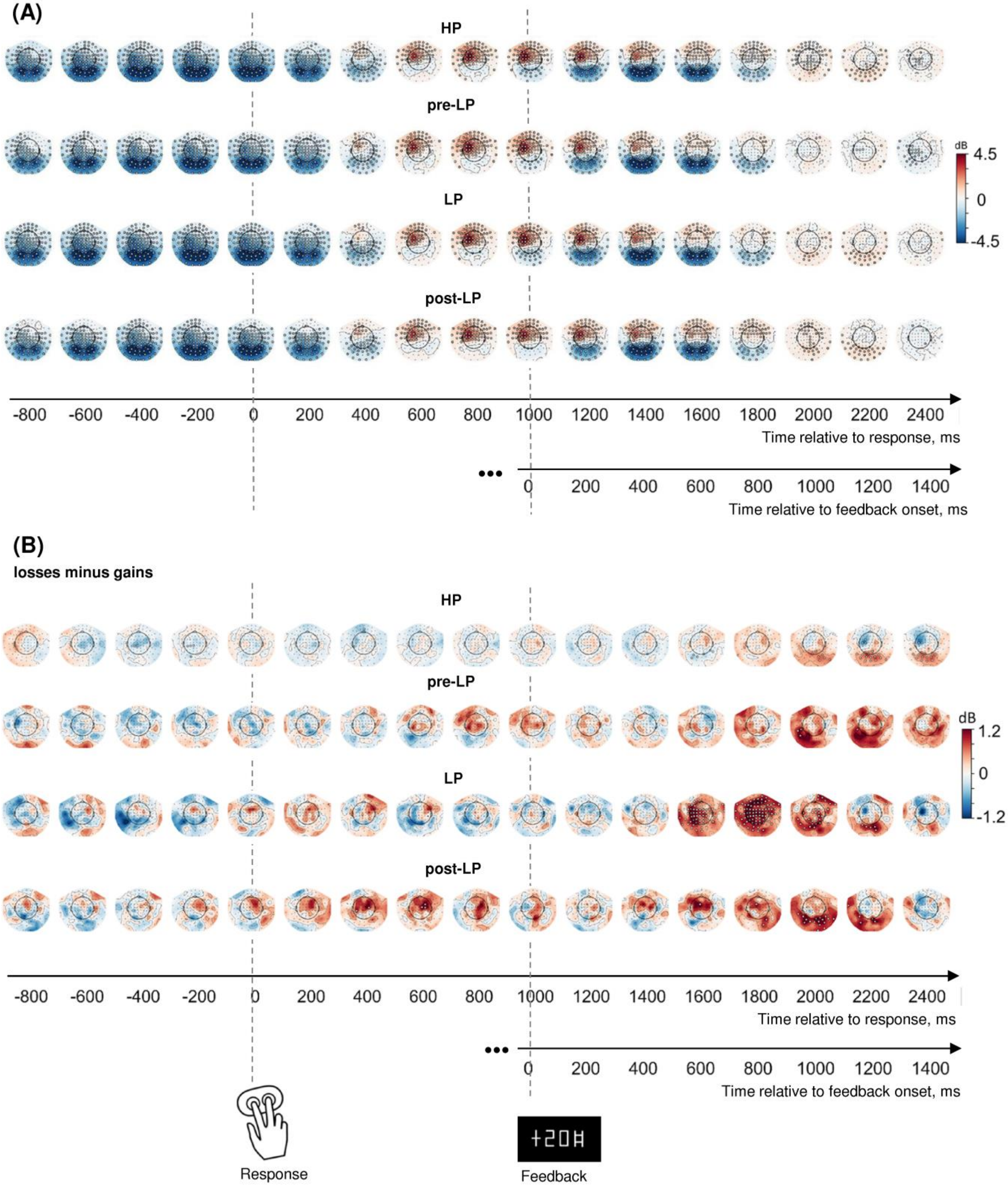

**Supplementary Figure S1**. Sequences of topographic maps of the power of β-band oscillations in response-locked data; significant sensors are indicated by open circles. Each map represents β power averaged over a 200-ms time window (-100 – 100 ms relative to the timestamps indicated).

**(A)** Each choice type minus baseline (t-statistics, uncorrected);

**(B)** 'Losses' minus 'gains', for each choice type (LMEM statistics, FDR-corrected for 17 time windows x 102 sensors).

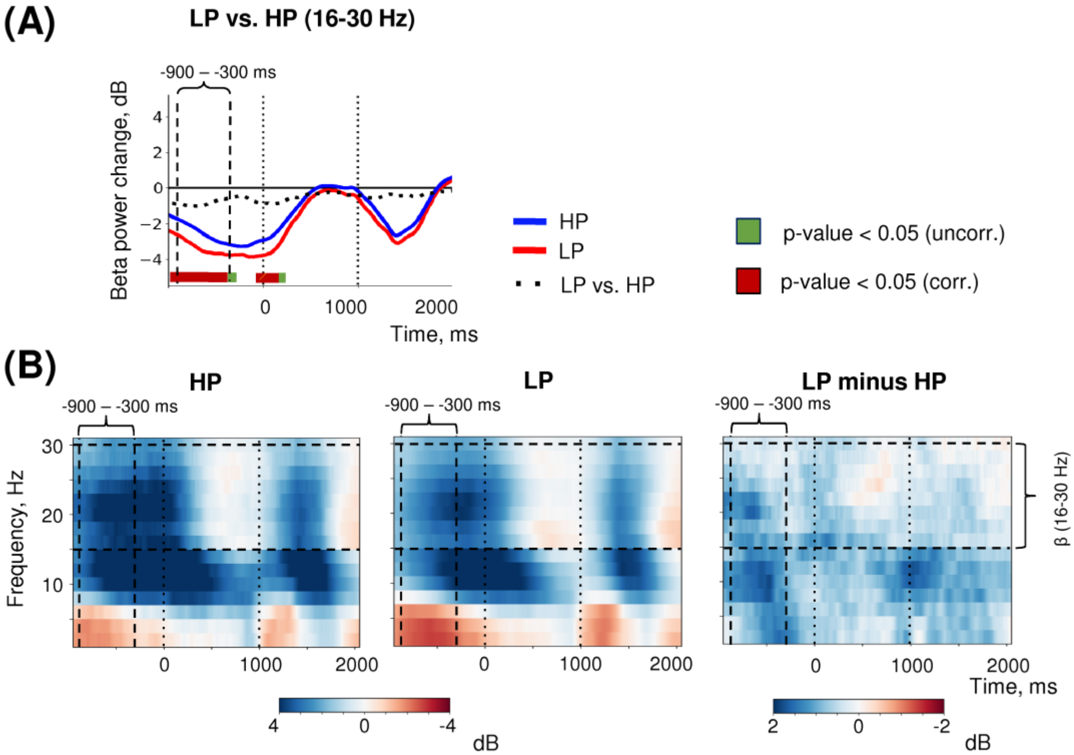


**Supplementary Figure S2**. Additional time-frequency analysis for the decision-related effect.

**(A)** Timecourses of β-band power averaged over three most significant sensors (shown in the inset at the top). Each graph represents two timecourses and the difference between them. Lines at the bottom of each graph indicate significance of the difference (t-test, $p < 0.05$; green: uncorrected; brown: FDR-corrected over time points).

**(B)** Event-related spectral perturbations of MEG power averaged within the same three sensors obtained in the main LMM analysis.

'0' on the timeline corresponds to response, and '1000' – to feedback onset.

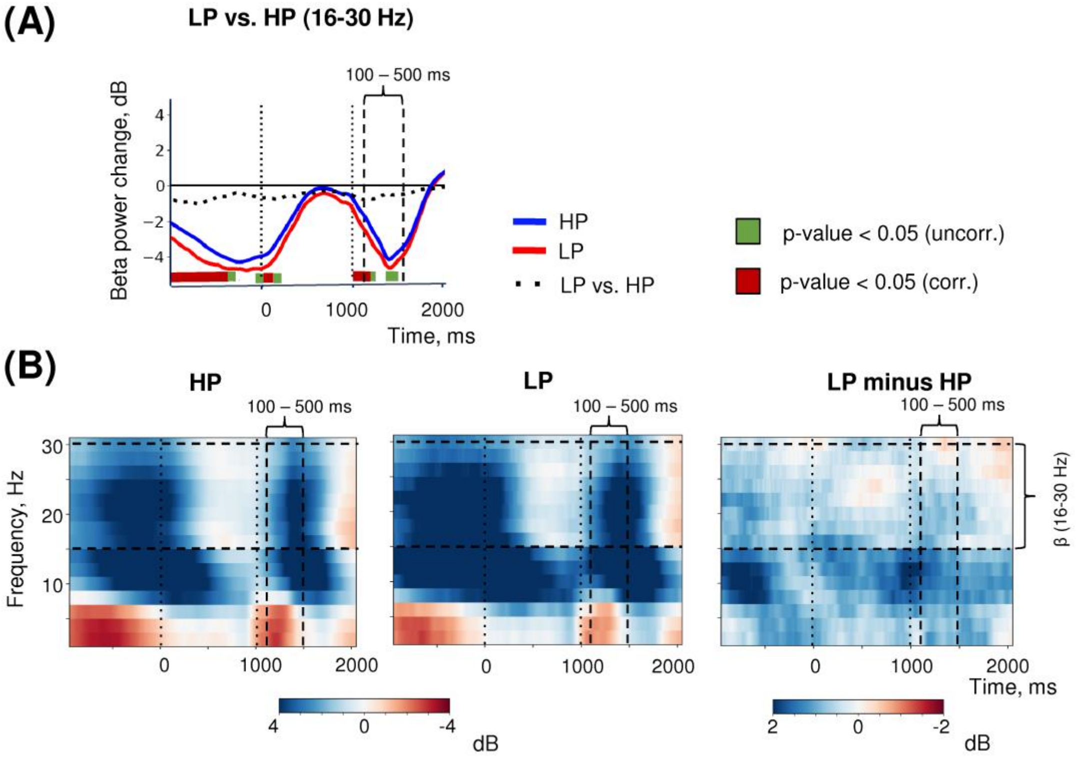


**Supplementary Figure S3**. Additional time-frequency analysis for the early feedback effect.

**(A)** Timecourses of β-band power averaged over three most significant sensors (shown in the inset at the top). Each graph represents two timecourses and the difference between them. Lines at the bottom of each graph indicate significance of the difference (t-test, $p < 0.05$; green: uncorrected; brown: FDR corrected over time points).

**(B)** Event-related spectral perturbations of MEG power averaged within the same three sensors obtained in the main LMM analysis.

'0' on the timeline corresponds to response, and '1000' – to feedback onset.

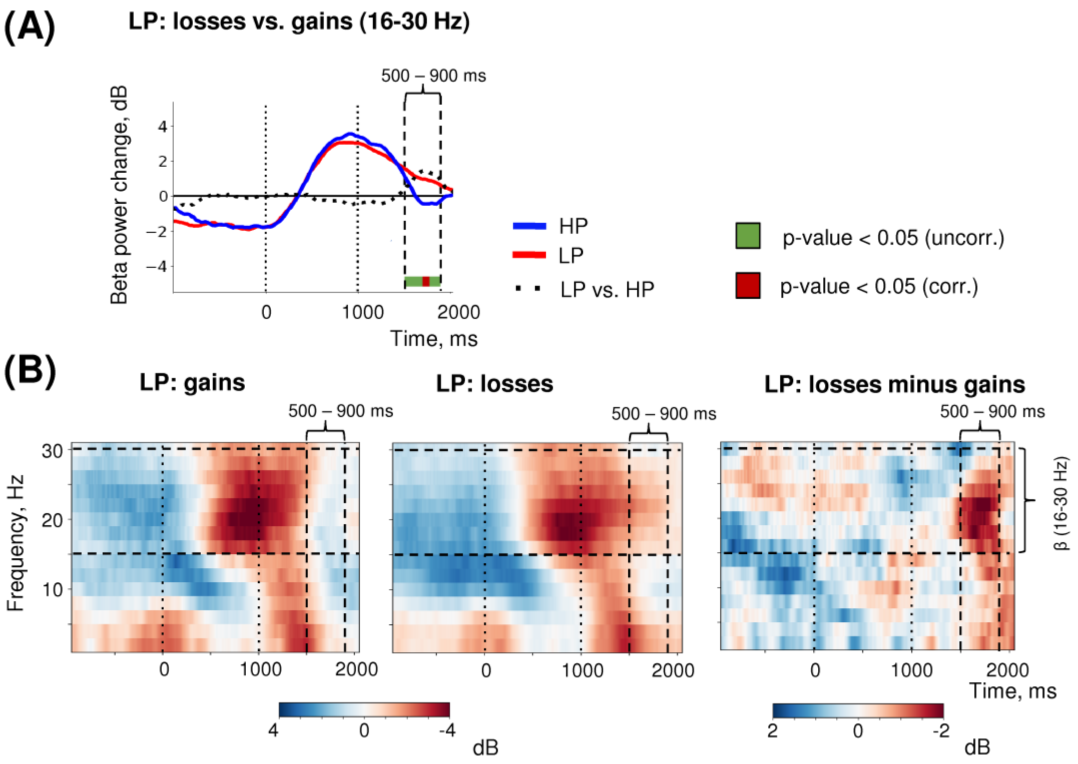


**Supplementary Figure S4**. Additional time-frequency analysis for the late feedback effect in the anterior cluster of sensors.

**(A)** Timecourses of β-band power averaged over three most significant sensors in the anterior cluster (shown in the inset at the top). Each graph represents two timecourses and the difference between them. Lines at the bottom of each graph indicate significance of the difference (t-test, $p < 0.05$; green: uncorrected; brown: FDR corrected over time points).

**(B)** Event-related spectral perturbations of MEG power averaged within the same three sensors obtained in the main LMM analysis.

'0' on the timeline corresponds to response, and '1000' – to feedback onset.

**Supplementary Table S1**. Choice Types and Overall Behavioral Statistics

| Choice type | Trials[¥]: Previous → **Current** → Next | Number of trials | % of trials | RT, ms (*M* ± *SD*) |
|---|---|---|---|---|
| HP | A → **A** → A | 3148 | 64.4 | 1383 ± 88 |
| pre-LP | A → **A** → DA | 600 | 12.3 | 1502 ± 96 |
| LP | A → **DA** → A | 539 | 11.0 | 1806 ± 95 |
| post-LP | DA →**A** → A | 602 | 12.3 | 1514 ± 97 |

[¥] A – advantageous choice, DA – disadvantageous choice.

**Supplementary Table S2**. LMM statistics for the response time

| | **df1** | **df2** | **F** | **p** |
|---|---|---|---|---|
| **Choice type** | **3** | **3519.35** | **17.09** | **<0.001***** |
| Previous feedback | 1 | 3519.89 | 1.85 | 0.174 |
| **Choice type x Previous feedback** | **3** | **3515.64** | **4.72** | **0.003**** |

** – p<0.01, *** – p<0.001.

**Supplementary Table S3**. LMM statistics for the effects of choice types, feedback valence, and their interaction for beta power change averaged across the pre-specified sensors and time windows based on the previous LMM analysis results[¥]

| Effect | Factor | df1 | df2 | F | p |
|---|---|---|---|---|---|
| Decision-related effect | **Choice type[†]** | **3** | **4324.4** | **28** | **<0.001*****|
| Early feedback-related effect | **Choice type[†]** | **3** | **4317.9** | **13.9** | **<0.001***** |
| | Feedback | 1 | 4316.5 | 0.02 | 0.874 |
| | Choice type x Feedback | 3 | 4316.9 | 1.65 | 0.176 |
| Late feedback-related effect (posterior cluster) | **Choice type** | **3** | **4318.6** | **2.8** | **0.037*** |
| | **Feedback** | **1** | **4316.8** | **13.7** | **<0.001***** |
| | **Choice type x Feedback[†]** | **3** | **4317.4** | **13.3** | **<0.001***** |
| Late feedback-related effect (anterior cluster) | **Choice type** | **3** | **4322.8** | **9.78** | **<0.001***** |
| | **Feedback** | **1** | **4316.9** | **37.3** | **<0.001***** |
| | **Choice type x Feedback[†]** | **3** | **4319.1** | **11** | **<0.001***** |

[¥] LMM statistics were computed to ensure that the effects obtained for beta power at the level of single sensors and time bins survived after averaging beta values across space and time and should not be considered in isolation from the previous results.

[†] Significance is a consequence of the procedure used to define time intervals and clusters of sensors (cf. Figure 2).

* – p<0.05, *** – p<0.001 (Tukey HSD).
